# Identification of novel autophagy inducers by accelerating lysosomal clustering against Parkinson’s disease

**DOI:** 10.1101/2023.11.07.566147

**Authors:** Yuki Date, Yukiko Sasazawa, Mitsuhiro Kitagawa, Kentaro Gejima, Ayami Suzuki, Hideyuki Saya, Masaya Imoto, Eisuke Itakura, Nobutaka Hattori, Shinji Saiki

**Affiliations:** Department of Biology, Graduate School of Science and Engineering, Chiba University, Inage-ku, Chiba 263-8522, Japan; Department of Neurology, Juntendo University Faculty of Medicine, Tokyo, Japan; Research Institute for Diseases of Old Age, Juntendo University Graduate School of Medicine, Tokyo, Japan; Division for Development of Autophagy Modulating Drugs, Juntendo University Faculty of Medicine, Tokyo, Japan; Division of Gene Regulation, Institute for Advanced Medical Research, School of Medicine, Keio University, Tokyo, Japan; Division of Gene Regulation, Cancer Center, Fujita Health University, Toyoake, Japan; Department of Biology, Graduate School of Science, Chiba University, Inage-ku, Chiba 263-8522, Japan; Neurodegenerative Disorders Collaborative Laboratory, RIKEN Center for Brain Science, Saitama, Japan; Department of Neurology, Institute of Medicine, University of Tsukuba, Ibaraki, Japan

## Abstract

Autophagy-lysosome pathway plays an indispensable role in the intracellular protein quality control system, degrading abnormal organelles and proteins. Among these proteins is α-Synuclein (αSyn), which is associated with the pathogenesis of Parkinson’s disease (PD). However, the activation of this lysosome-dependent degradation strategy is restricted by enzyme complementation. In this study, we focused on the phase of autophagosome-lysosome fusion around the microtubule organizing center (MTOC) that leads to αSyn degradation. Through high-throughput chemical screening, we identified six clinically available drugs that enhance autophagy and can accumulate lysosomes around the MTOC from approximately 1,200 drugs screened. We further demonstrated that these compounds induce lysosomal clustering through a JIP4-TRPML1-dependent mechanism, which is associated with autophagy induction. Among these, the lysosomal clustering compound albendazole was observed to promote the autophagy-dependent degradation of Triton-X-insoluble proteasome inhibitor-induced aggregates (p62). In a cellular PD model, albendazole boosted the degradation of insoluble αSyn, an effect that was reversed upon the addition of bafilomycin A1. Our results suggest that lysosomal clustering can facilitate the breakdown of protein aggregates. Therefore, compounds that promote lysosomal clustering may offer a promising therapeutic strategy against neurodegenerative diseases characterized by the presence of aggregate-prone proteins.

## Introduction

Parkinson’s disease (PD), the second most prevalent progressive neurodegenerative disorder after Alzheimer’s disease, is characterized by the depletion of dopamine due to the loss of dopaminergic neurons in the substantia nigra pars compacta. PD affects a broad range of both motor and non-motor functions and is observed in approximately 1% of individuals over the age of 65. One of the primary therapies for PD is dopamine supplementation, often using medications like levodopa. While levodopa has a transformative effect, its chronic use can result in adverse effects such as wearing-off and dyskinesia. Since current treatments don’t prevent the loss of dopaminergic neurons, there’s an urgent need for new therapeutic strategies to address the challenges of PD.

Recent pathological insights into PD indicate that the aggregation and deposition of the α-Synuclein (αSyn) protein, known as Lewy bodies, within dopaminergic neurons in the substantia nigra, play a critical role in PD’s etiology (Spillantini et al., 1997; Postuma et al., 2015). There’s increasing interest in stimulating autophagy—a primary protein degradation system—using small-molecule compounds as a therapeutic approach, especially due to its role in clearing αSyn aggregates (Webb et al., 2003; Gao et al., 2019; Choi et al., 2020a). During autophagy, an isolation membrane engulfs part of the cytoplasm, forming a double-membraned organelle known as an autophagosome. This organelle then fuses with lysosomes, creating autolysosomes where lysosomal hydrolases break down the vesicles’ contents. Significantly, rapamycin, a recognized autophagy inducer, has been shown to inhibit αSyn aggregation in in vivo models and improve motor functions (Crews et al., 2010; Dehay et al., 2010). Yet, even with the discovery of autophagy inducers like rapamycin, no fully effective treatment is available. For instance, while nilotinib boosts autophagic αSyn clearance in αSyn transgenic PD model mice, its effects in clinical trials were underwhelming. Therefore, there’s a pressing need for compounds that can dismantle aggregated proteins using innovative mechanisms.

Recent studies highlight the importance of lysosomal distribution in regulating autophagy and its associated genes. Lysosomes clustering around the microtubule organizing center (MTOC) play a key role in managing autophagic flux. They do this by facilitating autophagosome formation and inhibiting the mechanistic target of rapamycin complex 1 (mTORC1). The close physical association of these clustered lysosomes with autophagosomes eases their fusion (Kimura et al., 2008; Korolchuk et al., 2011a). The transport of lysosomes to the MTOC is dependent on the dynein/dynactin complex and encompasses several protein pathways: (i) the small GTPase Rab7-Rab7 effector Rab-interacting lysosomal protein (RILP) pathway (Johansson et al., 2007; Rocha et al., 2009), (ii) the transient receptor potential mucolipin 1 (TRPML1) apoptosis-linked gene 2 (ALG2) pathway (Li et al., 2016), and (iii) the lysosomal membrane protein TMEM55B-JNK-interacting protein 4 (JIP4) pathway (Willett et al., 2017).Furthermore, we recently discerned an additional pathway involving phosphorylated JIP4-TRPML1-ALG2 that’s induced by oxidative stress (Sasazawa et al., 2022). We also hypothesize that the phosphorylation status of JIP4 at T217 by CaMK2G serves as a determining switch, altering its binding affinity and thus dictating the pathway JIP4 employs.

Notably, disruptions in lysosome and autophagosome transport can contribute to neurodegenerative diseases. For example, dynactin mutations are linked to Perry’s disease, a rare hereditary neurodegenerative condition marked by autosomal dominant parkinsonism (Konno et al., 2017). Such mutations disrupt lysosomal distribution, leading to hindered autophagy and cell death (Ishikawa et al., 2014). Additionally, leucine-rich repeat kinase 2 (LRRK2), a causative protein in PD, is essential in regulating autophagosome and lysosomal trafficking, particularly in partnership with JIP4 (Bonet-Ponce et al., 2020; Kluss et al., 2022a; b).

On the other hand, mounting evidence suggests that protein aggregates, including those causing neurodegenerative diseases, gather around MTOCs and are degraded via the autophagy-lysosome pathway. Proteasome inhibitors, for example, aggregate near the nucleus and undergo autophagic degradation (Jänen et al., 2010; Choi et al., 2020b). In the same vein, the pathogenic protein mutant huntingtin, linked to Huntington’s disease, forms aggresomes near the nucleus and is degraded autophagically (Waelter et al., 2001; Ma et al., 2022). Similarly, the αSyn aggregates that form Lewy bodies in PD patients are found near the MTOC (Olanow et al., 2004), hinting at the possibility that MTOC-centric autophagy aids in their removal. Prior research has shown that overexpressing Arl8b hampers the degradation of αSyn-A53T (Korolchuk et al., 2011a), suggesting that lysosomal retrograde trafficking is vital in αSyn-A53T’s breakdown.

Given this, we hypothesize that compounds encouraging lysosomal clustering near MTOCs could minimize the distance between lysosomes, autophagosomes, and degradation substrates, thereby enhancing the degradation of protein aggregates. Guided by this idea, we’ve developed a screening system. We first employed a high-content image screening method to identify autophagy inducers, with a focus on lysosomal clustering. Consequently, we identified a novel set of autophagy inducers that encourage lysosomal clustering. Importantly, we demonstrated that these compounds effectively degrade αSyn aggregates, underscoring the potential of enhancing lysosomal clustering as a strategy to eliminate protein aggregates.

## Results

### Cell-based screening for compounds inducing lysosomal clustering

To identify compounds that enhance the degradation of αSyn aggregates, we initially screened for those that induce lysosomal clustering. We subsequently assessed their autophagy-inducing activity (Fig. 1A).

**Fig. 1.**
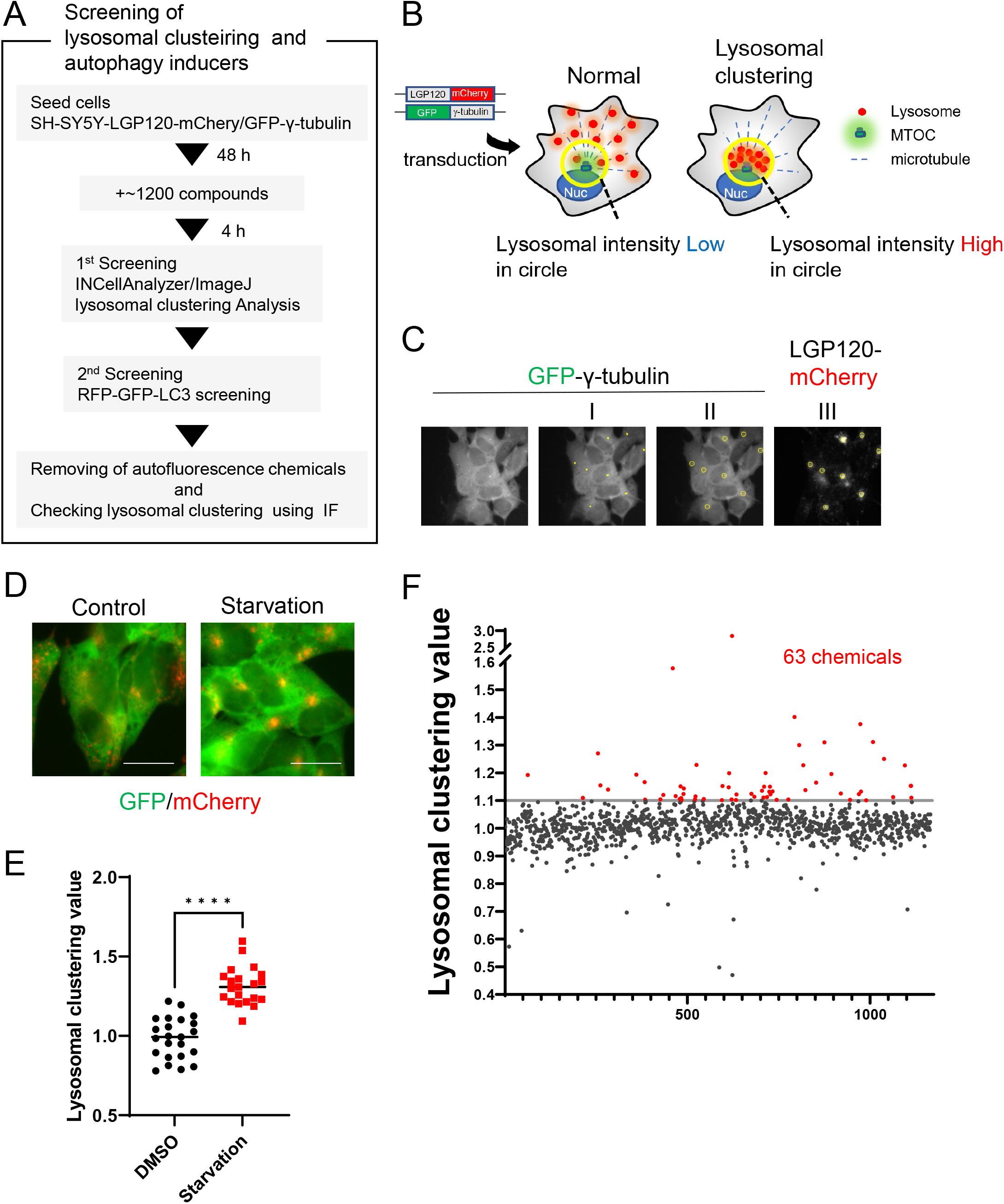
The method of lysosomal clustering chemicals screening. A. The scheme for lysosomal-clustering chemical screening. B. Strategies for screening lysosomal clustering compounds. If lysosomes accumulate around the MTOC, the fluorescent intensity (red) in the circle increases. C. INCellAnalyzer2200 images of SH-SY5Y cells co-expressing GFP-γ-tubulin and LGP120-mcherry. I-III detail the Image J lysosomal clustering analysis procedure. (I) From the γ-tubulin image, the MTOC position is extracted using Image J processing. (II) A circle about 7 μm in diameter is placed at the central coordinates of the MTOC position. (III) The circle is reflected onto the LGP120 image to measure the LGP120 fluorescence intensity (lysosomal clustering value) within the circle. D. SH-SY5Y cell lines co-expressing GFP-γ-tubulin and LGP120-mcherry were treated with either control medium or starvation medium (positive control). The results were analyzed using INCellAnalyzer2200 and ImageJ. E. The graph displays the lysosomal clustering value (see methods) (n > 20). **** P < 0.0001, Wilcoxon test. F. The graph demonstrates the fold change in the lysosome clustering value for 1200 chemicals relative to the control. Chemicals with over a 1.1-fold change in lysosome clustering value relative to the control were identified as lysosome clustering chemicals.

For the identification process, we devised a high-content imaging screening system utilizing INCellAnalyzer2200. As a starting point, we engineered SH-SY5Y cells to stably express both LGP120-mCherry (a lysosomal marker) and GFP-γ-tubulin (an MTOC marker) (Fig. 1B). Using the INCellAnalyzer2200, we captured fluorescence images (from both GFP and mCherry). The distribution of lysosomes was then quantified via the ImageJ software. We pinpointed the MTOC location using the GFP-γ-tubulin spot signal, then designated a roughly 7 µm circle in diameter centered on the MTOC. The intensity of the LGP120-mCherry signal within this circle was gauged, and its ratio compared to the cell’s overall intensity was determined (Fig. 1C). This ratio, when related to the control, was difined as the lysosomal clustering value.

To ensure the accuracy of our approach, we cultivated the cells under starvation conditions, using this as a positive control, and subsequently assessed lysosomal distribution. Starvation resulted in evident lysosomal clustering, noticeable in fluorescence images and corroborated by the heightened lysosomal clustering value (Fig. 1D, E). This underscores the utility of our high-content imaging screening system.

Leveraging this screening method, we assessed approximately 1200 compounds, all of which have received clinical approval in Japan. Of these, 63 compounds exhibited a high lysosomal clustering value (greater than 1.1) (Fig. 1F, TableS1).

### Topoisomerase inhibitor and benzimidazole induce lysosomal clustering and autophagy

Next, to identify the autophagy inducers among these compounds, we assessed the autophagic activity of these compounds using RFP-GFP tandem fluorescent-tagged LC3 (R-G-LC3) as a second screening (Fig. 2A) (Takayama et al., 2017). We established SH-SY5Y cells stably expressing R-G-LC3. During the fusion of autophagosomes with mammalian lysosomes in these cells, GFP fluorescence is quenched due to its pH sensitivity and is degraded by lysosomal proteases. In contrast, RFP is resistant to both acidic conditions and lysosomal proteases, resulting in accumulation in lysosomes. The distinct properties of RFP and GFP enabled us to evaluate autophagic activity sensitively by measuring the fluorescence ratio of RFP/GFP in a cell using flow cytometry (Fig. S1).

**Fig. 2.**
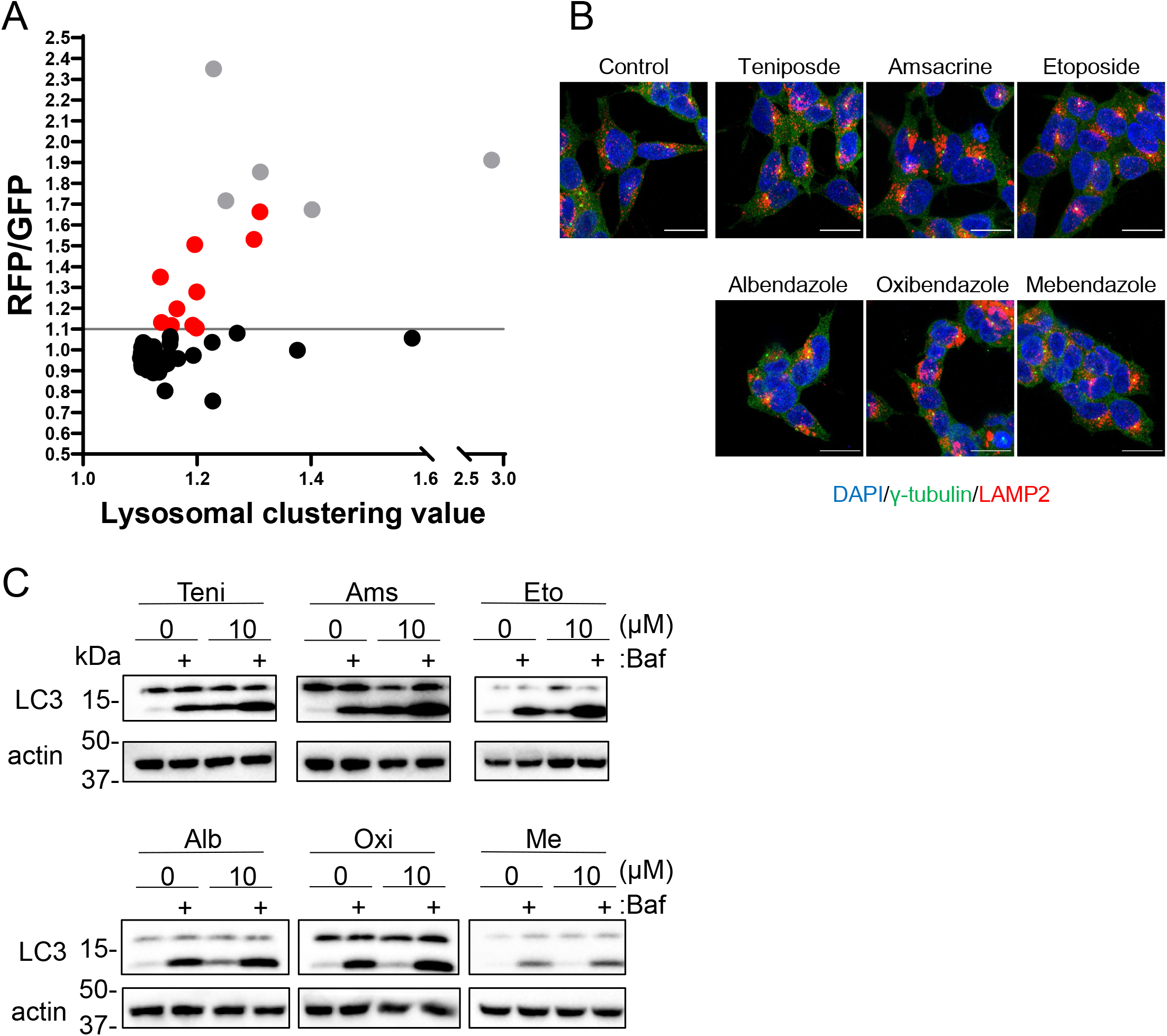
Identification of autophagy inducers via lysosomal clustering. A. SH-SY5Y cells expressing RFP-GFP-LC3 were analyzed by FACS after treatment with 63 lysosomal clustering chemicals for 24 h. The graph illustrates the fold change in the RFP/GFP ratio for these 63 chemicals relative to the control. Auto-fluorescent chemicals were excluded if they had > 1.1-fold change in RFP/GFP ratio relative to the control. After confirming endogenous lysosome accumulation using IF, six lysosomal clustering autophagy inducers were ultimately identified. Gray dots represent autofluorescence chemical data. B. SH-SY5Y cells were treated with teniposide (10 μM), amsacrine (10 μM), etoposide (10 μM), albendazole (10 μM), oxibendazole (1 μM), and mebendazole (5 μM) for 4 h. Cells were then fixed and stained with the specified antibody and DAPI. C. SH-SY5Y cells underwent treatment with lysosome clustering chemicals for 6 h. Cell lysates were immunoblotted with the indicated antibodies.

From the secondary screening of the 63 hit compounds identified in the first screening, we pinpointed 15 compounds that induce autophagy with lysosomal clustering (Fig. 2A, TableS1). As five of these compounds are auto-fluorescent chemicals (denoted by gray dots in Fig. 2A), we further evaluated the effects of the remaining 10 compounds on endogenous lysosomal clustering. We eventually identified 6 compounds that lead to lysosomal clustering and subsequent enhancement of autophagy (Fig. 2B). These compounds were categorized into two types: the topoisomerase II inhibitors (topo-i), including teniposide, amsacrine, and etoposide; and the benzimidazole class anthelmintics, comprising albendazole, oxibendazole, and mebendazole. An immunofluorescence assay of R-G-LC3-expressing SH-SY5Y cells revealed that both starvation and treatment with either teniposide or albendazole promote autophagosomal perinuclear clustering and increase their co-localization with lysosomes (autolysosomes) (Fig. S2).

Furthermore, an autophagy flux assay conducted via western blot indicated that these compounds exhibit LC3B lipidation, an effect that was amplified by the lysosomal inhibitor bafilomycin A1 (Mizushima and Yoshimori, 2007) (Fig. 2C). Drawing from the results, we have successfully identified a novel class of six autophagy inducers that promote lysosomal clustering.

### Six hit compounds induce lysosomal clustering and autophagy in a mTORC1-independent manner

Lysosomal retrograde transport governs autophagic flux by facilitating the formation of autophagosomes. This is achieved through the inhibition of mTORC1 and by aiding in the fusion between autophagosomes and lysosomes (Kimura et al., 2008; Korolchuk et al., 2011a). Consequently, we examined whether the six compounds modulate mTORC1 activity through the regulation of lysosomal positioning. This was done by monitoring the phosphorylation states of mTOR and the mTORC1 downstream effectors, such as ribosomal S6 protein kinase (p70S6K), ribosomal S6 protein (S6), and ULK1 (Klionsky et al., 2016). Torin1, a selective mTOR inhibitor, along with starvation medium, was observed to hinder the phosphorylation of S6, p70S6K, mTOR, and ULK1. In contrast, the six compounds only marginally reduced the levels of p-p70S6K without influencing p-S6, p-mTOR, and p-ULK1 (Fig. 3A). Given the lack of a clear link between lysosomal accumulation and mTORC1 activity, as demonstrated by Torin1 treatment (Fig. 3B, C, Fig. S2), the six compounds seem to induce autophagy primarily independent of mTORC1 inhibition.

**Fig. 3.**
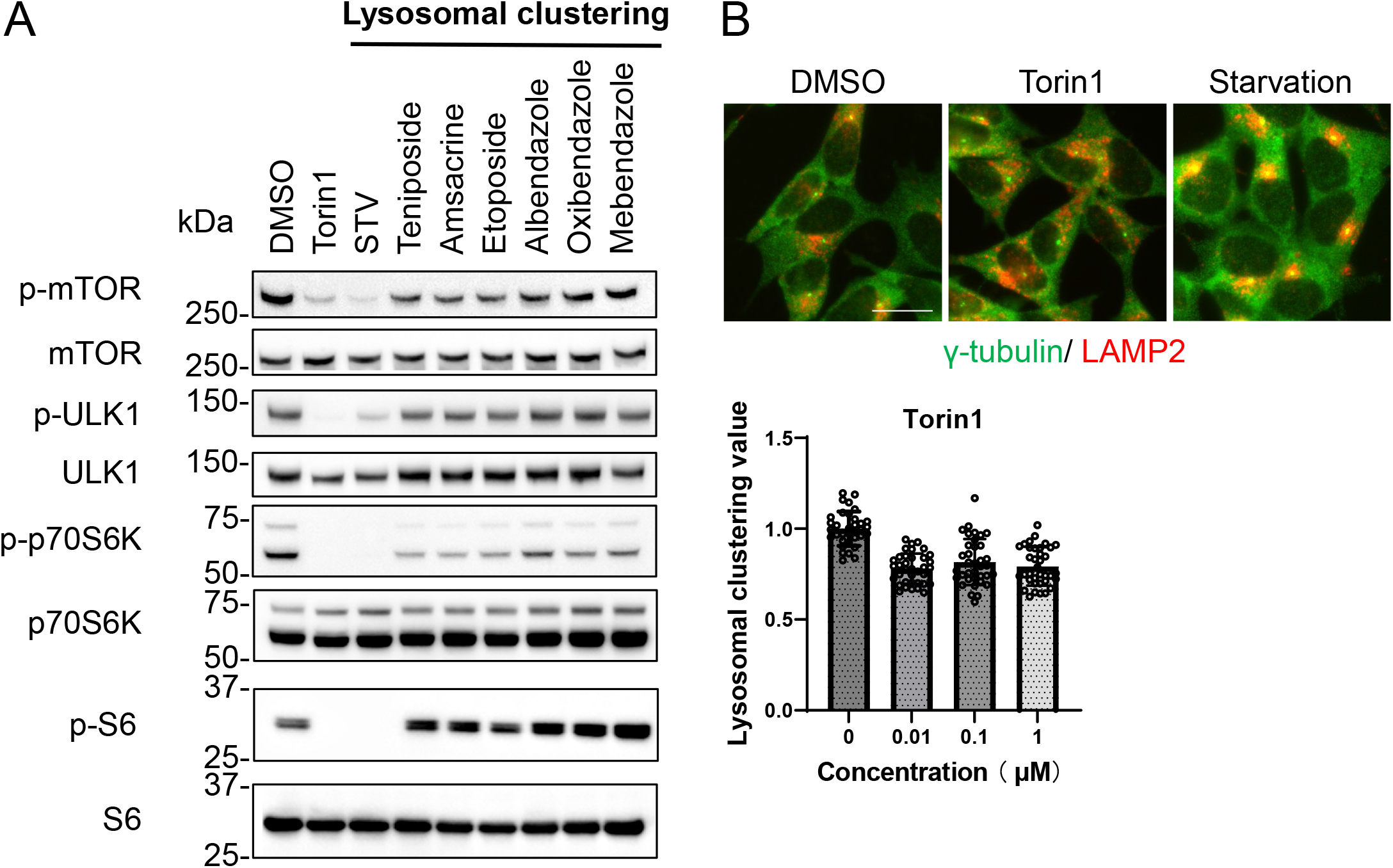
Lysosomal clustering chemicals are not dependent on mTORC1 activity. A. SH-SY5Y cells were treated with starvation medium, torin1 (1 μM), teniposide (10 μM), amsacrine (10 μM), etoposide (10 μM), albendazole (10 μM), oxibendazole (1 μM), or mebendazole (5 μM) for 4 h. Cell lysates were then immunoblotted with the specified antibody. B. SH-SY5Y cells received treatment with torin1 (1 μM) for 4 h. These cells were fixed, stained with anti-γ-tubulin (green) and anti-LAMP2 antibody (red), and imaged using INCellAnalyzer2200. Scale bar represents 20 μm. INCellAnalyzer2200 images were then analyzed using ImageJ for lysosomal clustering. The graph illustrates the lysosomal clustering value (n > 30). Data is presented as means ± SD.

### Topoisomerase inhibitor and benzimidazole transport lysosomes dependently through the JIP4-TRPML1 pathway

Various factors are known to regulate lysosomal clustering. Four primary dynein-mediated pathways for lysosomal clustering have been identified: Rab7-RILP (Johansson et al., 2007), TRPML1-ALG2 (Li et al., 2016), TMEM55B-JIP4 (Willett et al., 2017), and TRPML1-JIP4-ALG2 (Sasazawa et al., 2022) pathways. To discern which pathways are implicated in the lysosomal retrograde transport initiated by these hit compounds, we first examined the effects of Rab7, RILP, ALG2, TRPML1, TMEM55B, and JIP4 knockdown on the lysosomal distribution spurred by teniposide and albendazole. These compounds typify topoisomerase inhibitors (topo-i) and benzimidazole class anthelmintics, respectively. The efficiency of knockdown against these genes was validated via western blotting and quantitative reverse transcription–PCR (Fig. S3). JIP4 and TRPML1 knockdown hindered teniposide-induced lysosomal clustering, while TMEM55B, ALG2, Rab7, and RILP knockdown did not. Conversely, albendazole-induced lysosomal clustering was stymied by JIP4, TRPML1, ALG2, and Rab7 knockdown, but TMEM55B and RILP knockdown had no such effect (Fig. 4A, B). These observations suggest that topo-i-driven lysosomal clustering is mediated by JIP4 and TRPML1, while albendazole-induced clustering involves JIP4, TRPML1, ALG2, and Rab7. Furthermore, in JIP4 knockout (KO) cells, lysosomal clustering induction for all six chemicals was entirely nullified (Fig. 4C, D, Fig. S4). Immunostaining using JIP4 antibodies revealed increased co-localization of JIP4 with LAMP2 under both teniposide and albendazole treatment conditions (Fig. 4E). Collectively, these results suggest that while teniposide and albendazole may stimulate lysosomal clustering via different pathways, both TRPML1 and JIP4 are instrumental in each.

**Fig. 4.**
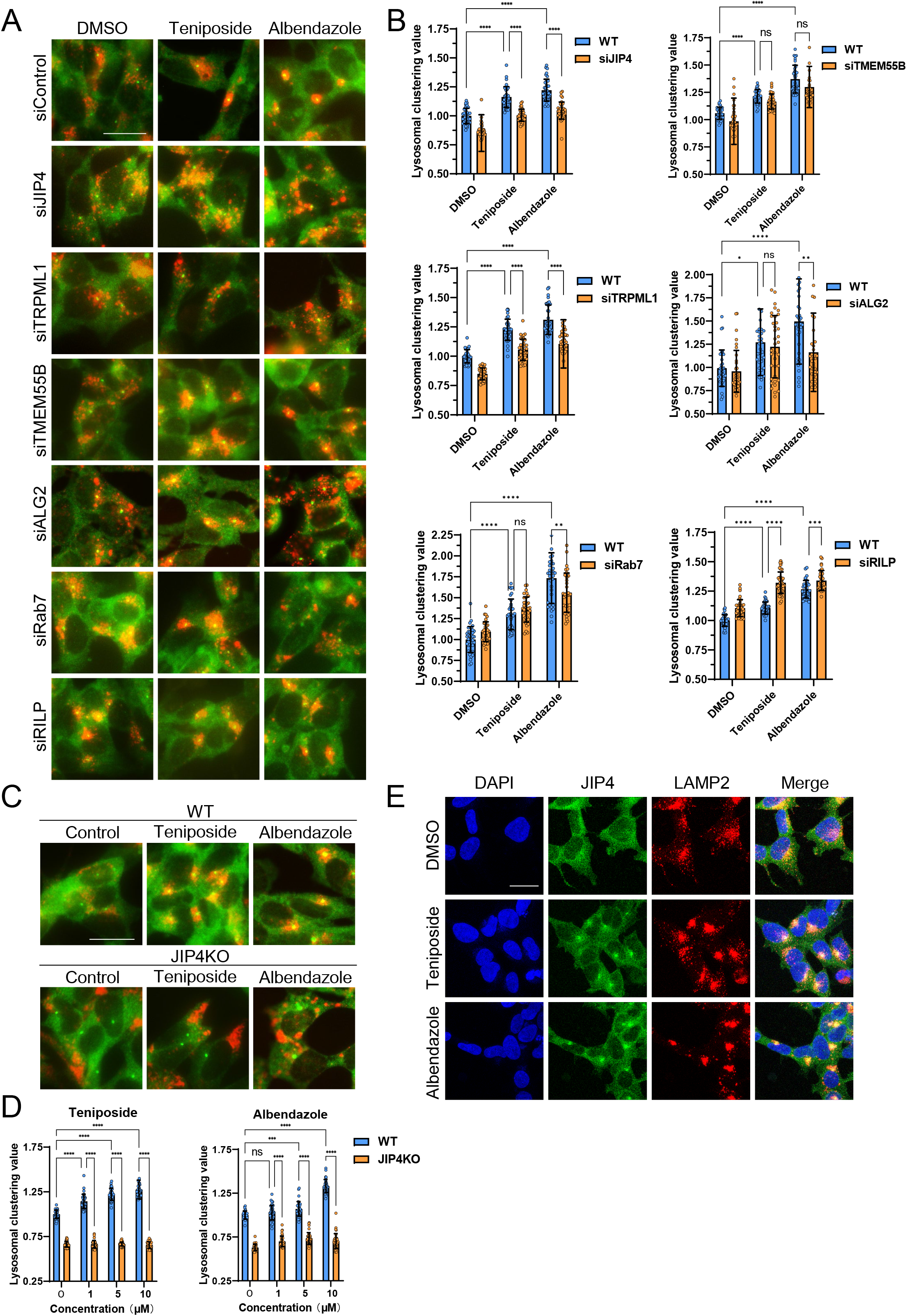
Lysosome clustering chemicals accumulate lysosomes in MTOC in a JIP4-dependent manner. A. SH-SY5Y cells were transfected with the indicated siRNAs for 48 h and then treated with teniposide (10 μM) and albendazole (10 μM) for 4 h. Cells were fixed and stained with an anti-γ-tubulin (green) and an anti-LAMP2 antibody (red). Images were captured using INCellAnalyzer2200. Scale bar: 20 μm. B. INCellAnalyzer2200 images were analyzed using ImageJ for lysosomal clustering. The graph displays the lysosomal clustering value (n > 30). Data is presented as means ± SD. ****P < 0.0001, ***P < 0.001, **P < 0.01, *P < 0.05; N.S., not statistically different; two-way ANOVA and Tukey’s test. The experiment was replicated at least three times. C. SH-SY5Y WT and JIP4 knockout cells were cultured in 96-well black plates and treated with teniposide (1, 5, or 10 μM) and albendazole (1, 5, or 10 μM). Cells were fixed and stained with an anti-γ-tubulin (green) and an anti-LAMP2 antibody (red). Images were captured using INCellAnalyzer2200. D. INCellAnalyzer2200 images were analyzed using ImageJ for lysosomal clustering. The graph displays the lysosomal clustering value for WT or JIP4KO cells (n > 30). Data is presented as means ± SD. ****P < 0.0001, ***P < 0.001, **P < 0.01, *P < 0.05; N.S., not statistically different; two-way ANOVA and Tukey’s test. The experiment was replicated at least three times. E. SH-SY5Y cells were treated with teniposide (10 μM) and albendazole (10 μM) for 4 h. Cells were fixed and stained with anti-JIP4 (green) and anti-LAMP2 (red) antibodies, as well as DAPI.

### Topoisomerase inhibitors drive lysosomal transport via phosphorylated JIP4, unlike benzimidazole

We previously determined that JIP4 plays a crucial role in regulating lysosomal clustering triggered by both oxidative stress and starvation. Interestingly, these two processes employ distinct pathways influenced by the phosphorylation status of JIP4. Specifically, oxidative stress results in the phosphorylation of JIP4 at T217 through CaMK2G activation, steering lysosomal retrograde transport in tandem with TRPML1 and ALG2. In contrast, starvation mediates lysosomal retrograde transport predominantly through the TRPML1-ALG2 and TMEM55B-JIP4 pathways, without JIP4 phosphorylation (Sasazawa et al., 2022).

Given this background, we sought to ascertain whether lysosomal clustering induced by topoisomerase inhibitors (topo-i) and benzimidazole hinges on JIP4 phosphorylation. To investigate this, we employed Jak3 inhibitor VI, previously identified as an inhibitor of the JIP4 kinase CaMK2G, which oversees oxidative stress-induced lysosomal clustering (Sasazawa et al., 2022). Notably, the Jak3 inhibitor VI effectively nullified lysosomal clustering prompted by the three topo-i compounds. In contrast, benzimidazole remained largely unaffected (Fig. 5A, B). Moreover, phostag-PAGE analyses revealed that all three topo-i compounds lead to JIP4 phosphorylation, an effect counteracted by Jak3 inhibitor VI (Fig. 5C). Further, siCaMK2G inhibited lysosomal clustering induced by all three topo-i compounds (Fig. 5D, E). This suggests that CaMK2G-phosphorylated JIP4 is pivotal for both topo-i-induced lysosomal clustering and oxidative stress response. Additionally, rescue experiments in JIP4 KO SH-SY5Y cells, which re-express either flag-tagged wild-type JIP4 or a phosphorylation-defective variant (T217A JIP4), revealed that only the cells with wild-type flag-JIP4 could recover the teniposide-induced lysosomal clustering phenotype (Fig. 5F). This underscores the indispensability of JIP4 phosphorylation at T217 for the changes in lysosomal distribution induced by topo-i. In summary, while topo-i-induced lysosomal clustering is orchestrated by phosphorylated JIP4 (T217) in conjunction with TRPML1 (but not ALG2), benzimidazole-driven lysosomal retrograde transport, exemplified by albendazole, engages the TRPML1-JIP4-ALG2 and Rab7 pathways.

**Fig. 5.**
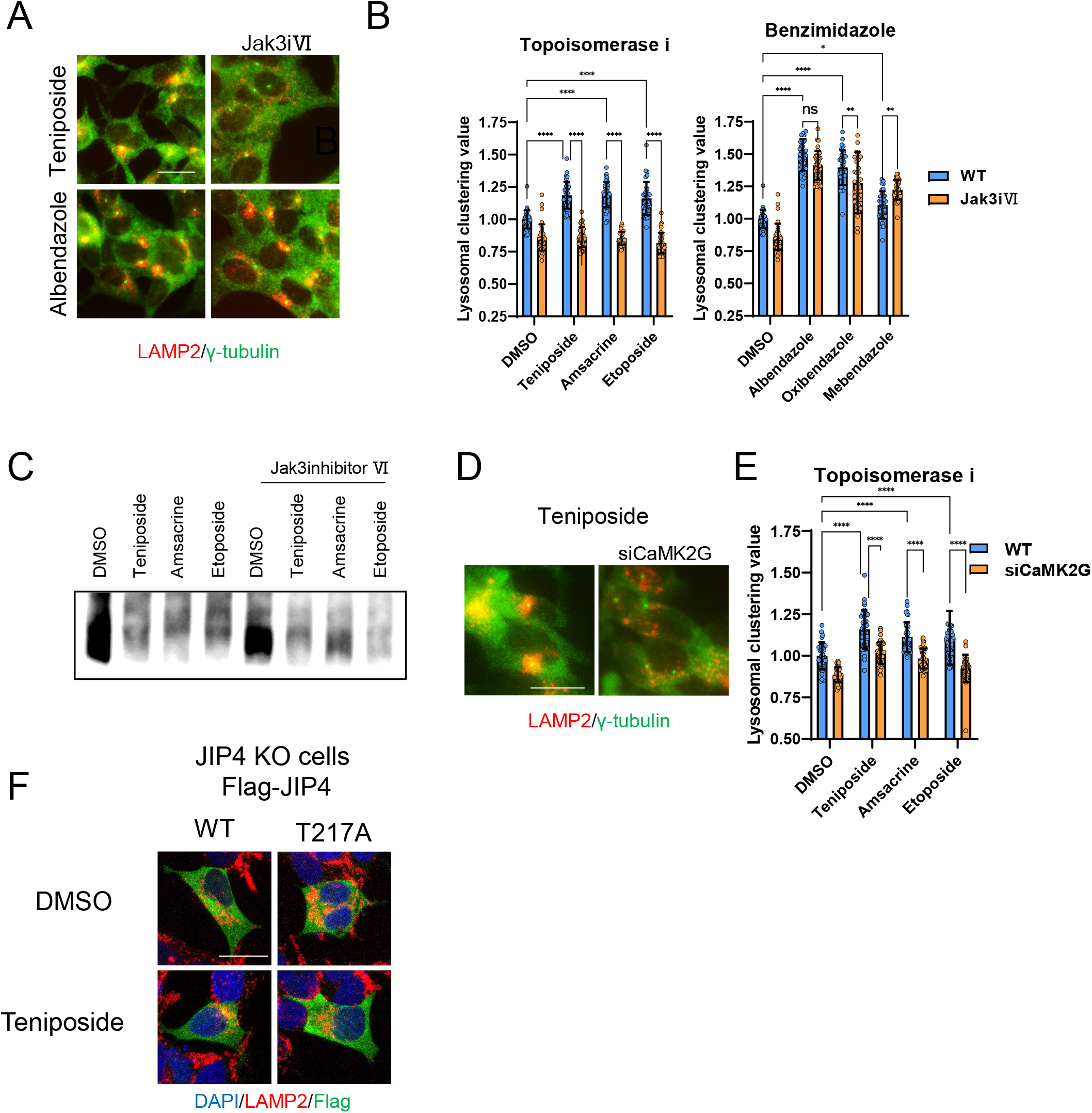
Lysosomal clustering by topoisomerase inhibitor requires phosphorylation of JIP4. A. SH-SY5Y cells were treated with DMSO, teniposide (10 μM), amsacrine (10 μM), etoposide (10 μM), albendazole (10 μM), oxibendazole (1 μM), mebendazole (5 μM) with or without Jak3 inhibitor VI for 4 h. Cells were fixed and stained with an anti-γ-tubulin (green) and an anti-LAMP2 antibody (red). Images were captured using INCellAnalyzer2200. B. INCellAnalyzer2200 images were analyzed using ImageJ for lysosomal clustering. The graph displays the lysosomal clustering value for DMSO or Jak3 inhibitor VI (n > 30). Data is presented as means ± SD. ****P < 0.0001, ***P < 0.001, **P < 0.01, *P < 0.05; N.S., not statistically different; two-way ANOVA and Tukey’s test. The experiment was replicated at least three times. C. SH-SY5Y cells were treated with teniposide (10 μM), amsacrine (10 μM), etoposide (10 μM) with or without Jak3 inhibitor VI for 4 h. Cell lysates underwent Phos-tag PAGE and were immunoblotted with an anti-JIP4 antibody. D. SH-SY5Y cells were transfected with CaMK2G siRNAs for 48 h and then treated with teniposide (10 μM), amsacrine (10 μM), etoposide (10 μM) for 4 h. Cells were fixed and stained with an anti-γ-tubulin (green) and an anti-LAMP2 antibody (red). Images were captured using INCellAnalyzer2200. Scale bar: 20 μm. E. INCellAnalyzer2200 images were analyzed using ImageJ for lysosomal clustering. The graph displays the lysosomal clustering value (n > 30). Data is presented as means ± SD. ****P < 0.0001, two-way ANOVA and Tukey’s test. The experiment was replicated at least three times. F. JIP4 KO cells were transfected with flag-tagged JIP4 (WT and T217A) for 24 h and treated with teniposide (10 μM) for 4 h. Cells were fixed and stained with the indicated antibodies. Scale bar: 20 μm.

### These compounds transport autophagosomes and lysosomes in a JIP4-dependent manner, thereby inducing efficient autophagy

Next, we investigated whether lysosomal clustering is essential for the autophagy triggered by these lysosomal clustering compounds. To confirm this, we utilized a quantifiable HaloTag-LC3B assay to measure autophagic flux (Yim et al., 2022). In cells expressing HaloTag-LC3B, the HaloTag degrades within lysosomes in the absence of ligands. However, when ligands are introduced, the HaloTag portion accumulates inside lysosomes. This accumulation serves as an estimate of autophagic activity (Yim et al., 2022). We expressed HaloTag-LC3B in both parental SH-SY5Y cells and JIP4 KO cells and gauged autophagic flux by adding topo-i and benzimidazole. Consequently, JIP4 KO cells displayed reduced accumulation of the cleaved Halo bands in the in-gel fluorescence images for both compounds (Fig. 6A, B). These findings reveal that both topo-i and benzimidazole provoke JIP4-dependent lysosomal clustering, subsequently inducing autophagy.

**Fig. 6.**
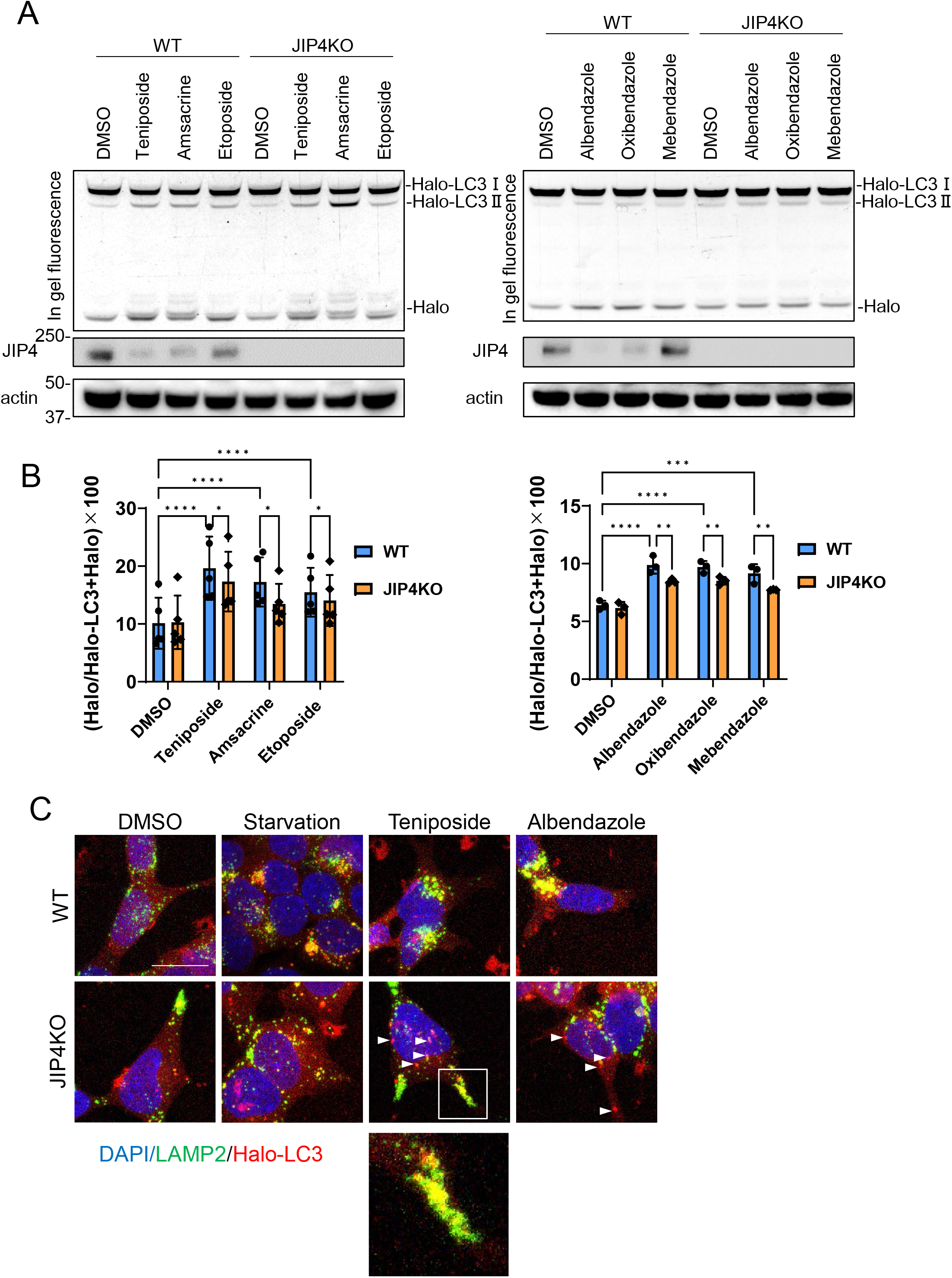
Lysosomal clustering dependent on JIP4 is slightly involved in autophagy activity. A. SH-SY5Y cells stably expressing Halo-LC3 were labeled for 20 min with 100 nM of tetramethyl rhodamine (TMR)-conjugated ligand in a nutrient-rich medium. After washing with PBS and incubating the cells in normal medium for 30 min, cells were treated with DMSO, teniposide (10 μM), amsacrine (10 μM), etoposide (10 μM), albendazole (10 μM), oxibendazole (1 μM), and mebendazole (5 μM) for 4 h. Cell lysates were immunoblotted with the indicated antibody and analyzed by in-gel fluorescence detection. B. Quantification of results shown in (A). HaloTMR band intensity was normalized by the sum of the band intensities HaloTMR-LC3B and HaloTMR. The vertical axis of the graph represents the intensity multiplied by 100. Mean values of data from five or three experiments are shown. The data are presented as the means ± SD. ****P < 0.0001, ***P < 0.001, **P < 0.01, *P < 0.05, two-way ANOVA and Tukey’s test. C. Cells treated as in A were fixed and stained with an anti-LAMP2 antibody (green). These cells were imaged by a confocal microscope. Scale bar, 20 μm. White arrows indicate Halo-LC3 dots. The magnified image shows LAMP2 (green) and Halo dots (red) accumulating at the cell periphery.

Furthermore, an immunofluorescence assay demonstrated that starvation, teniposide, and albendazole treatments promote autophagosomal perinuclear clustering and augment their co-localization with lysosomes, forming autolysosomes (Fig. 6C). Intriguingly, in JIP4 KO cells, lysosomal clustering was inhibited as was the accumulation of autophagosomes to the MTOC. Specifically, for DMSO and teniposide treatments, the accumulation of autolysosomes (indicated by the colocalization of Halo-LC3 with lysosomes) was observed at the cell periphery following teniposide treatments (Fig. 6C, enlarged image), suggesting the induction of autophagy. As indicated in our previous study (Ishikawa et al., 2019), inhibiting retrograde vesicular transport (for example, through dynactin knockdown) causes both autophagosomes and lysosomes to disperse to the cell periphery, resulting in residual autolysosome formation there, preserving some autophagic flux. This is likely why a slight yet notable decline in autophagic flux was observed in JIP4 KO cells treated with topo-i and benzimidazole. On the other hand, in the presence of albendazole or under starvation conditions in JIP4 KO cells, the accumulation of autolysosomes at the cell periphery was infrequently observed. This supports the notion that lysosomal clustering induced by albendazole, or starvation was also mediated by pathways other than JIP4, such as the Rab7-mediated pathway.

Additionally, autophagosomes that did not co-localize with lysosomes were detected in the cytoplasm following treatments with teniposide, albendazole, and under starvation conditions in JIP4 KO cells (Fig. 6C, indicated by a white arrow). These findings imply that teniposide and albendazole-induced autophagy necessitates JIP4-dependent lysosomal movement to induce autophagy and to facilitate the fusion of autophagosomes with lysosomes. Moreover, these results suggest JIP4’s potential role not only in lysosomal retrograde transport but also in autophagosomal retrograde transport. In summary, in JIP4 KO cells, lysosomes and autophagosomes are directed to the cell periphery, retaining some level of autophagic flux.

### Albendazole can degrade proteasome inhibitor MG132-induced aggregates

Because topo-i exhibits cytotoxicity and is clinically used as an anti-cancer drug, we next explored how albendazole might induce the degradation of protein aggregates through autophagy. Previous reports have already indicated that aggregates caused by proteasomal inhibition can be degraded by autophagy (Jänen et al., 2010; Choi et al., 2020), and the significance of lysosomal clustering has also been highlighted (Zaarur et al., 2014). Indeed, MG132, a well-known proteasomal inhibitor, increased the levels of p62 and poly-ubiquitinated proteins in the Triton-X insoluble fraction. This increase declined in a time-dependent manner after the washout of MG132 (Fig. 7A). Furthermore, the reduction in p62 and ubiquitin levels, following the MG132 washout, was markedly suppressed in autophagy-deficient FIP200 KO cells (Fig. 7B). These results suggest that basal autophagy degrades the protein aggregates induced by MG132. Notably, when treated with albendazole in conjunction with the MG132 washout, p62 reduction occurred more rapidly than with DMSO alone (Fig. 7C). The effects of albendazole, however, were suppressed in FIP200 KO cells (Fig. 7D). These findings imply that albendazole can degrade protein aggregates formed by a proteasome inhibitor in an autophagy-dependent manner.

**Fig. 7.**
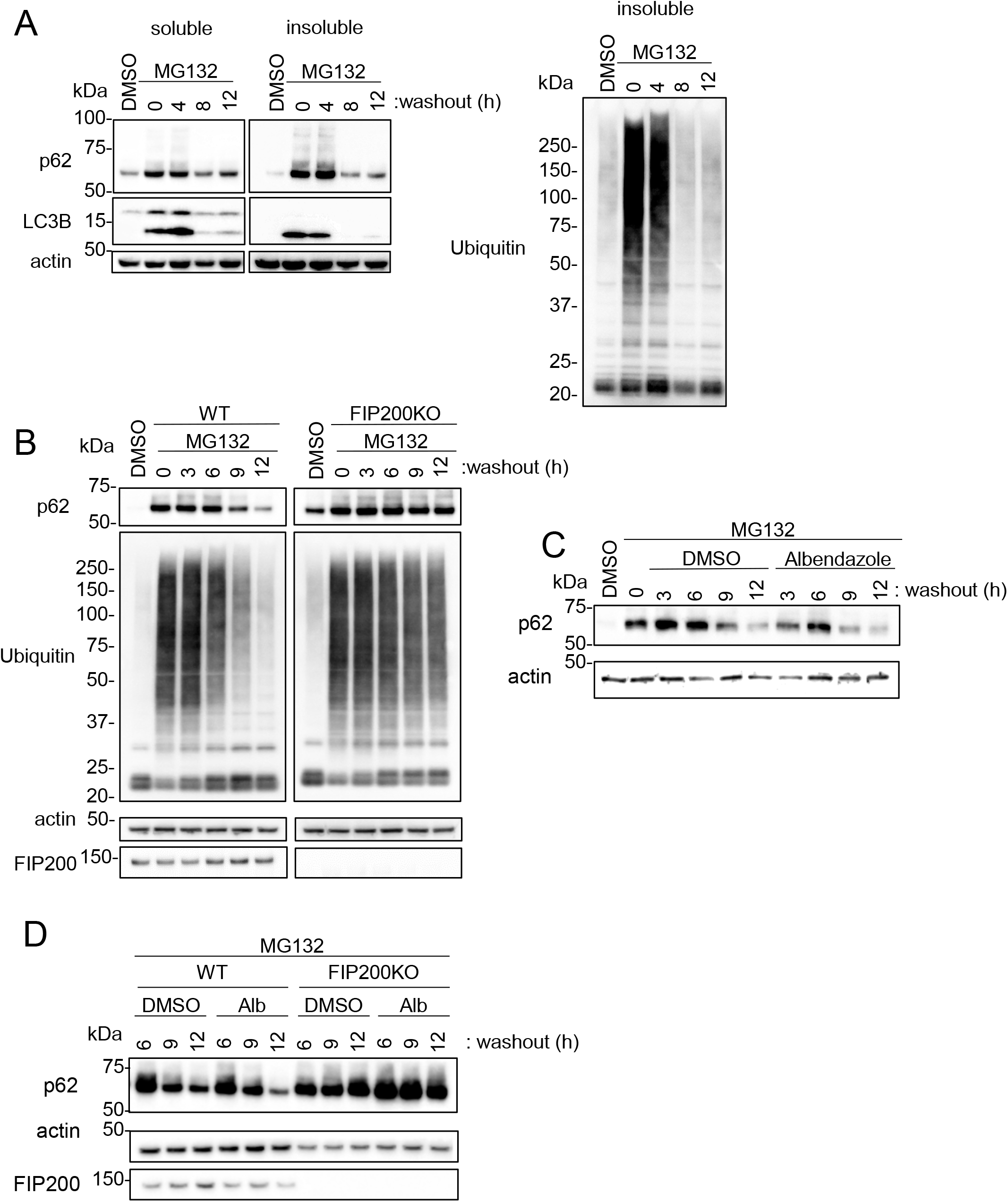
JIP4-dependent lysosomal clustering is involved in aggresome clearance by MG132. A. SH-SY5Y cells were treated with MG132 (1 μM) for 16 h to induce aggresome formation. After washing out MG132 with normal medium for intervals of 4, 8, or 12 h, cell lysates were separated into Triton X-100–soluble (soluble) and pellet fractions (insoluble), then subjected to SDS/PAGE/immunoblotting with the indicated antibody. B. WT or FIP200KO SH-SY5Y cells were treated with MG132 (1 μM) for 16 h to induce aggresome formation. After washing out MG132 with normal medium for intervals of 3, 6, 9, or 12 h, the same analysis as in A was performed. C. SH-SY5Y cells were treated with MG132 (1 μM) for 16 h. After washing out MG132 with either DMSO or albendazole in normal medium for intervals of 3, 6, 9, or 12 h, the same analysis as in A was performed. D. WT or FIP200KO SH-SY5Y cells were treated with MG132 (1 μM) for 16 h to induce aggresome formation. After washing out MG132 with DMSO or albendazole in normal medium for intervals of 6, 9, or 12 h, the same analysis as in A was performed.

### Lysosomal clustering is required for the degradation of α-Synuclein aggregate

Next, we explored whether albendazole can degrade the PD causative protein, αSyn. For this purpose, we used cellular models of PD wherein αSyn aggregation is prompted by the transduction of αSyn fibrils as seeds in αSyn overexpressing cells (Nonaka et al., 2010). These αSyn fibrils were introduced into the SH-SY5Y cell line, which stably expresses the green GFP-fused human αSyn. Notably, confocal microscopy analyses showed that upon the introduction of αSyn fibrils, lysosomes gathered around the formed αSyn aggregates (Fig. 8A). Albendazole further facilitated the concentration of lysosomes on these αSyn-GFP aggregates (Fig. 8B). Under electron microscopy (EM), the presence of albendazole increased the recruitment of autophagosomes and lysosomes to the αSyn aggregates compared to the control (Fig. 8C). These observations suggest that lysosomes actively converge to aid in breaking down αSyn aggregates.

**Fig. 8.**
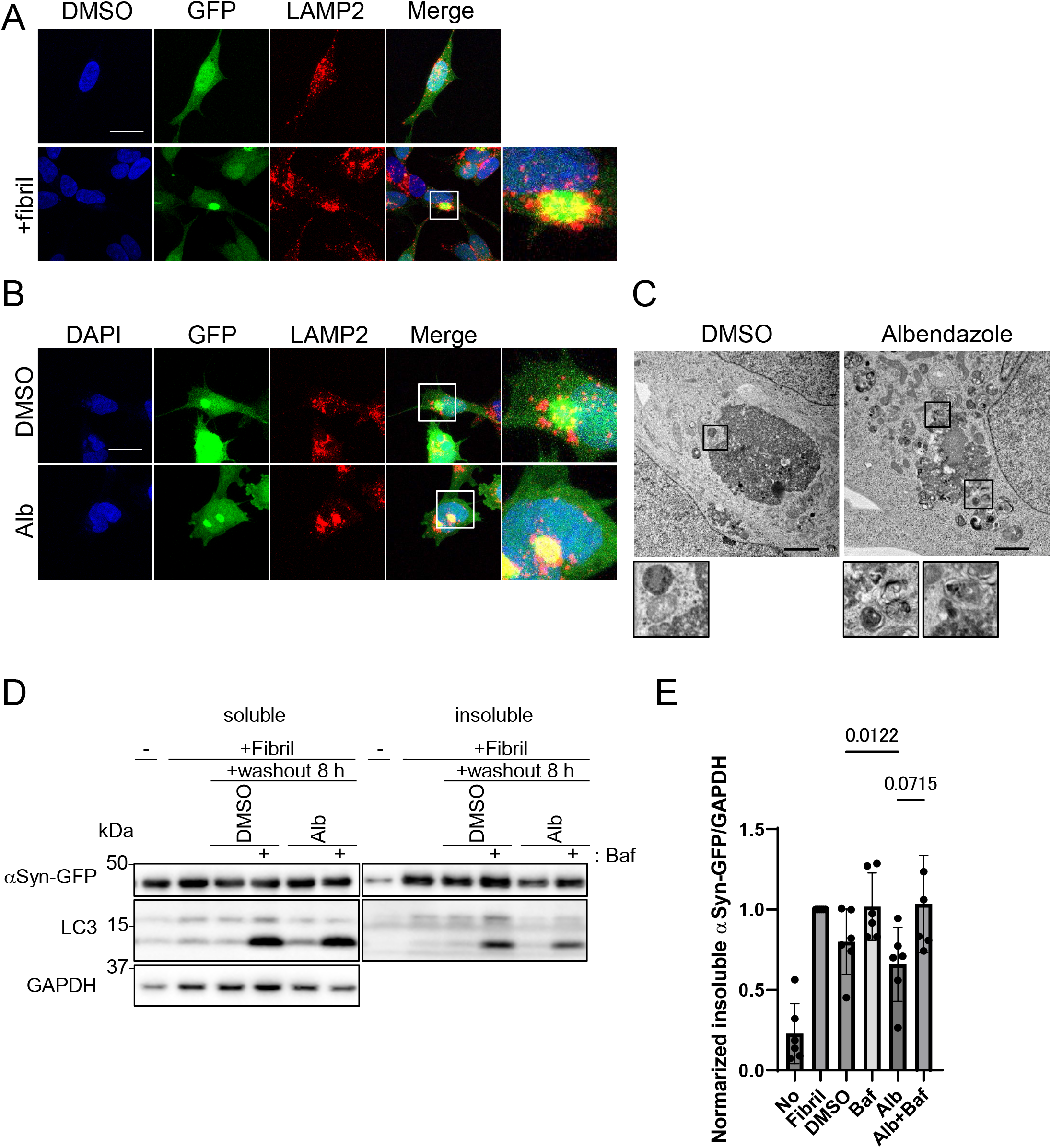
Albendazole reduces αSyn-GFP aggregates in a lysosome-dependent manner. A. SH-SY5Y cells overexpressing αSyn-GFP were transfected with αSyn fibril (0.2 µg/ml) using Lipofectamine3000. After 48 h, the cells were fixed and stained with an anti-LAMP2 antibody (red) and DAPI (blue). These cells were imaged by a confocal microscope. Scale bar, 20 μm. B. SH-SY5Y cells overexpressing αSyn-GFP were transfected with αSyn fibril (0.2 µg/ml) using Lipofectamine3000. After 48 h, and washing out the transfection reagent, SH-SY5Y cells were treated with DMSO or albendazole (10μM) with or without bafilomycin A1 (30 nM) for 8 h. Cells were fixed and stained with an anti-LAMP2 antibody (red) and DAPI (blue). These cells were imaged by a confocal microscope. Scale bar, 20 μm. C. SH-SY5Y cells overexpressing αSyn-GFP were transfected with αSyn fibril (0.2µg/ml) using Lipofectamine3000. After 48 h, and washing out the transfection reagent, SH-SY5Y cells were treated with DMSO or albendazole (10 μM), followed by EM analysis. Lysosome-like structures are enlarged at the bottom. Scale bar, 2 μm. D. SH-SY5Y cells overexpressing αSyn-GFP were transfected with αSyn fibril (0.2µg/ml) using Lipofectamine 3000. After 48 h, and washing out the transfection reagent, SH-SY5Y cells were treated with DMSO or albendazole (10 μM) with or without bafilomycin A1 (30nM) for 8 h. Cell lysates were separated into Triton X-100–soluble (soluble) and pellet fractions (insoluble), then subjected to SDS/PAGE/immunoblotting with the indicated antibody. E. The bar graph shows the insoluble αSyn-GFP ratio from D. The data are presented as the means ± SD. One-way ANOVA and Dunnett’s test.

To verify the degradation of αSyn aggregates by albendazole, we introduced αSyn fibrils to form the αSyn aggregates, washed out the fibrils, and then treated cells with albendazole, either with or without bafilomycin A1. Subsequently, we assessed the amount of αSyn in the Triton-X insoluble fraction (Fig. 8D). The introduction of αSyn fibrils notably increased αSyn-GFP levels, pointing to aggregated αSyn in the Triton-X insoluble fraction. However, these levels decreased post-fibril washout after an 8-hour culture with standard media. This decline was mildly obstructed by bafilomycin A1, suggesting that the basal level of autophagy degrades aggregated αSyn. Furthermore, treatment with albendazole showed a marked inclination to decrease αSyn-GFP levels compared to the DMSO treatment, and this decrease was also hindered by bafilomycin A1 (Fig. 8D, E). This pattern indicates that albendazole mediates the degradation of αSyn aggregates through autophagy. Taken together, our results suggest that albendazole enhances the autophagy-mediated degradation of αSyn-GFP aggregates.

## Discussion

In this study, we developed a novel autophagy-inducer screening method focusing on lysosomal clustering and assessed if these compounds could degrade αSyn. We created a high-throughput screening system to quantify lysosomal accumulation by measuring lysosomes located within a circular region centered on the MTOC. This was achieved through image analysis with INCellAnalyzer2200 and ImageJ Fiji (Fig. 1). We discovered that topo-i and benzimidazole promote lysosomal clustering via JIP4-TRPML1 (Fig. 2,4). Specifically, topo-i induces lysosomal clustering through the phosphorylation of JIP4 at T217 (Fig. 5). Compounds that cause lysosomal clustering may facilitate the fusion of autophagosomes and lysosomes near the MTOC by also transporting autophagosomes via JIP4 (Fig. 6). Additionally, we observed that albendazole induces lysosomal accumulation in αSyn aggregates located around the nucleus, subsequently degrading these aggregates in a lysosome-dependent manner (Fig. 8,9). Our results indicate that lysosomal clustering plays a significant role in αSyn degradation. Thus, novel lysosomal clustering chemicals have potential as clinical drugs for Parkinson’s disease, targeting αSyn aggregation.

The mechanisms through which topo-i and benzimidazole induce lysosomal clustering are distinct. Our findings reveal that topo-i-driven lysosomal transport is regulated by JIP4 and TRPML1, but not ALG2. This transport relies on the phosphorylation of JIP4 at T217 by CaMK2G. However, a previous study found that oxidative stress-induced lysosomal retrograde transport is influenced by phosphorylation-JIP4-TRPML1-ALG2 (Sasazawa et al., 2022). Therefore, topo-i prompts lysosomal clustering via a unique mechanism, involving phosphoJIP4 and TRPML1 without ALG2.

Earlier research has highlighted that JIP4 serves as a scaffold protein. Specifically, JIP4 binds to p38 MAPK, aiding its activation and undergoing phosphorylation in the process (Kelkar et al., 2005; Pinder et al., 2015). Another study reported that curcumin-induced lysosomal clustering was inhibited with p38 inhibitors (Willett et al., 2017). Indeed, a p38 inhibitor prevented topo-i-induced lysosomal clustering, but it had no effect on benzimidazole (Fig. S5A, B). Western blot analysis also showed that topo-i triggered p38 phosphorylation (Fig. S5C). These data hint at topo-i-induced lysosomal clustering’s potential link with p38. Since etoposide-induced DNA damage results in p38 phosphorylation (Khedri et al., 2019), it’s conceivable that topo-i-caused DNA damage might activate p38, facilitating lysosomal clustering.

Benzimidazole-induced lysosomal clustering is observed to be suppressed upon the knockdown of factors such as JIP4, TRPML1, Rab7, and ALG2. This suggests their potential involvement in benzimidazole’s retrograde lysosomal transport. Benzimidazole is known to bind to β-tubulin, disrupting microtubule-based processes in parasites (Lacey, 1988). We observed that oxibendazole, even in low concentrations, promoted lysosomal clustering. However, at high concentrations, it disrupted tubulin, inhibiting lysosomal clustering (Fig. S5D). This suggests that while intense tubulin polymerization conditions might disrupt tubulin, mild tubulin stimulation can lead to lysosomal accumulation. Thus, benzimidazole likely induces lysosomal clustering through a gentle stimulation of tubulin.

Lysosomal clustering within cells has diverse roles. It’s linked with the involvement of the PD-related protein LRRK2 in JIP4-mediated lysosomal positioning (Boecker et al., 2021; Bonet-Ponce et al., 2020) and plays a part in oxidative stress response (Sasazawa et al., 2022), pH regulation, and various cellular functions.

Our research showed that lysosomal clustering supports the fusion of autophagosomes with lysosomes, assisting in breaking down aggregates and other cellular components. Earlier studies reported that lysosomal retrograde transport enhances the fusion of autophagosomes and lysosomes during nutrient starvation, thus triggering autophagy (Korolchuk et al., 2011b). Consistent with these findings, our study discovered that in JIP4 knockout (JIP4KO) cells, Halo-LC3 degradation was hampered, implying a significant role for JIP4 in this process.

Ubiquitinated protein, p62 aggregation, and the proteasome itself, when blocked by the proteasome inhibitor MG132, are eventually degraded via the autophagy-lysosome pathway (Choi et al., 2020b; Jänen et al., 2010). It’s also been shown that forcing lysosomes to the cell periphery by overexpressing Arl8b inhibits the breakdown of aggregates induced by MG132 (Zaarur et al., 2014). Our research provides the first evidence for lysosomal clustering’s involvement in the degradation of not only MG132-induced aggresomes but also the αSyn fibril process. Our results highlight lysosomal clustering’s critical role in enhancing protein aggregate degradation.

Aggregates formed by the mutant huntingtin protein, associated with Huntington’s disease, are typically found both in the nucleus and the cytoplasm. These aggregates are known to attract autophagosomes via the autophagy adapter CCT2, leading to their subsequent autophagy-mediated degradation (Ma et al., 2022). During this process, the recruitment of lysosomes to the aggregates is evident. In our studies, we too observed an increase in lysosomal clustering and elevated LC3-II levels in the insoluble fraction during αSyn aggregate formation (Fig. 8D). This observation suggests that LC3-II might bind to αSyn aggregates, and the subsequent accumulation of autophagosomes and lysosomes around the aggregates could stimulate autophagy, leading to αSyn aggregate degradation. It’s possible that receptor proteins may link LC3-II to αSyn and the degradation of huntingtin protein aggregates.

In conclusion, our results emphasize that lysosomal clustering is pivotal in aggregate degradation. Albendazole, by enhancing autophagy through lysosomal accumulation, holds promise in aiding the breakdown of αSyn aggregates in Lewy body disease.

## Material & Methods

### Reagents

A chemical library consisting of about 1200 medicines clinically approved in Japan was supplied by Prof. Hideyuki Saya (Fujita Medical University, Japan). Teniposide, amsacrine, etoposide, albendazole, oxibendazole, and mebendazole were purchased from TCI in Japan. Bafilomycin A1 was purchased from Sigma-Aldrich (St. Louis, MO). Jak3 inhibitor Ⅵ was purchased from Merck Millipore (Burlington, MA). SB203580 was purchased from AdipoGen Life Science (Switzerland, Liestal, Epalinges).

### Cell culture

SH-SY5Y cells (American Type Culture Collection, ATCC#CRL-2266) were cultured in Dulbecco’s modified Eagle’s medium (Nacalai Tesque, Kyoto, Japan) supplemented with 10% fetal bovine serum (FBS; MP bio, Ringmer, UK), 100 U/ml penicillin/streptomycin (Nacalai Tesque, Kyoto, Japan), MEM non-essential amino acid solution (Thermo Fisher Scientific, Waltham, MA), 1 mM sodium pyruvate, and 2 mM L-glutamine at 37°C with 5% CO2. For starvation treatment, cells were washed with PBS and incubated in amino acid-free DMEM without serum (starvation medium (Wako)). Tetracycline-on (Tet-on) cells were generated by lentiviral transduction with a pCW57.1 vector (Addgene plasmid 41393, David Root lab) containing a single-vector Tet-on component and were cultured in the presence of 1 µg/ml doxycycline (Clontech, Mountain View, CA, USA) during induction.

### Stable cell line generation

LGP120-mCherry/GFP-γ-tubulin stably expressing cells were established by transfecting the respective vectors into SH-SY5Y cells using Lipofectamine LTX (Thermo Fisher Scientific), followed by selection with G418 (Roche Diagnostics, Indianapolis, IN). mCherry- and GFP-positive cells were sorted using a Cell Sorter Aria (BD) and plated on a 96-well plate. mCherry-GFP-LC3 and Halo-LC3 stably expressing SH-SY5Y cells were generated by either lentiviral or retroviral transduction. HEK293 cells were transiently co-transfected with lentiviral vectors using PEI MAX reagent (Polysciences, Warrington, PA, USA). Four hours post-transfection, the medium was replaced with fresh culture medium. After 72 hours of culturing, the growth medium containing the lentivirus was collected. SH-SY5Y cells were then incubated with the collected virus-containing medium for 48 hours. Uninfected cells were removed using 1 µg/ml puromycin or 5 µg/ml blasticidin S (Wako).

### Lysosomal clustering analysis

WT SH-SY5Y cells or those expressing GFP-γ-tubulin/LGP120-mCherry were cultured in 96-well black plates (Corning: 3603 or Greiner: 655892). After 48 hours of culture, cells were treated with compounds for 4 hours. Cells were subsequently fixed with 4% paraformaldehyde (Nakarai Tesque) and stained with Hoechst33342 (Invitrogen, H3570) for 30 minutes. Image capture of cells was performed using the High-content imager, INCellAnalyzer2200 (Cytiva). Lysosome distribution was quantified using Fiji software (Image J. ver.2.1.0/1.53c; Schindelin et al., 2012). The image analysis was as follows:

> Ⅰ: The MTOC position is identified from the γ-tubulin image using Image J processing (”ApplyLUT”⇒”Subtract Background”⇒”Threshold”).
>
> Ⅱ: A circle with a diameter of approximately 7 μm is centered on the MTOC coordinates.
>
> Ⅲ: This circle is superimposed on the LGP120 image, and the LGP120 fluorescence intensity within the circle is measured. The ratio of this fluorescence intensity to the whole-cell LAMP2 intensity is then calculated.

Processes Ⅰ-Ⅲ can be automatically analyzed using ImageJ programming. The lysosomal clustering value is defined as the ratio relative to the control.

### Immunoblotting

Western blot analysis was performed as previously described with minor modifications (Sasazawa et al., 2012, 2015, 2022). Cells were washed with cold PBS and lysed in lysis buffer (25 mM Tris–HCl pH 7.6, 150 mM NaCl, 1% NP-40, 1% sodium deoxycholate, 0.1% sodium dodecyl sulfate, and protease inhibitor cocktail) for 15 min on ice. The lysates were centrifuged at 20,000 g for 15 min to obtain soluble cell lysates. For SDS/polyacrylamide gel electrophoresis (SDS/PAGE), samples were mixed with 4×SDS sample buffer and boiled at 95°C for 5 min. 10μg of protein was added to each lane of the gel, separated by SDS-PAGE (Bio-Rad), and then transferred to a polyvinylidene difluoride (PVDF) membrane (Bio-Rad) using transfer buffer [25 mM Tris, 192 mM glycine, and 10% (vol/vol) methanol]. Immunoblot analysis was conducted with the indicated antibodies, and immunoreactive proteins were visualized using the West Dura Extended Duration Substrate (Thermo Fisher Scientific). The primary antibodies used were as follows: anti-LC3B (Cell Signaling Technology Inc., Danvers, MA, AB_2137707), anti-β-actin (Merck Millipore, AB_2223041), anti-phospho-mTORC1 (Ser2448) (Cell Signaling Technology Inc. AB_330970), anti-mTORC1 (Cell Signaling Technology Inc. AB_330978), anti-phospho-ULK1(Ser757) (Cell Signaling Technology Inc. AB_2665508), anti-ULK1 (Cell Signaling Technology Inc. AB_11178668), anti-phospho-p70S6K (Cell Signaling Technology Inc. AB_2269803), anti-p70S6K (Cell Signaling Technology Inc. AB_390722), anti-phospho-S6 (Cell Signaling Technology Inc., AB_916156), anti-S6 (Cell Signaling Technology Inc. AB_331355), anti-p62 (Abcam, AB_945626), anti-multi-ubiquitin (MBL, AB_592937), anti-FIP200 (D10D11) (Cell Signaling Technology Inc. AB_2797913), anti-GAPDH (Cell Signaling Technology Inc. AB_10622025), anti-TMEM55B (Proteintech, AB_2879391), anti-ALG2 (R&D Systems, Inc., AB_10972311), anti-JIP4 (Abcam, AB_299021), anti-GFP (Proteintech, AB_11182611).

To analyze the phosphorylated form of JIP4, lysates underwent 6% Phos-tag (50 μmol/l) acrylamide gel (FUJIFILM Wako Pure Chemical) electrophoresis. After electrophoresis, the gel was soaked in running buffer with 10 mM EDTA twice for 10 min each, and then in transfer buffer for 10 min. Proteins were transferred to a PVDF membrane and probed with the anti-JIP4 antibody.

### Flow cytometry

Cells detached with trypsin-EDTA were resuspended in 10% fetal bovine serum (FBS) and 1 µg/ml DAPI in PBS. They were then passed through a 70 µm cell strainer and analyzed using lasers equipped with NUV 375 nm (DAPI), 488 nm (GFP), and 561 nm (RFP) (BD). Dead cells were identified by DAPI staining. For each sample, 10,000 cells were acquired, and RFP/GFP fluorescence ratios (red fluorescence intensity divided by green fluorescence intensity) were calculated for RFP-positive cells.

### Immunofluorescence

Cells were cultured on coverslips and fixed with 4% paraformaldehyde (Nakarai Tesque) for 30 min. For immunostaining, fixed cells were permeabilized with 50 µg/ml digitonin in PBS for 15 min, blocked with 4% bovine serum albumin in PBS for 30 min, and then incubated with primary and secondary antibodies for 1 h. Fluorescence images were captured using an LSM880 confocal laser scanning microscope (Carl Zeiss, Oberkochen, Germany). The primary antibodies used were as follows: anti-LAMP2 (Development Studies Hybridoma Bank, Iowa City, IA; clone H4B4, AB_2134755), anti-γ-tubulin (abcam, AB_2904198) and anti-JIP4 (Thermo Fisher Scientific., AB_2642850). AB_2687580) and anti-p62 (abcam AB_56416).

### α-Synuclein aggregation assays

SH-SY5Y cells overexpressing α-Synuclein (αSyn)-GFP were cultured for 24 h. Cells were transfected with αSyn Fibril (0.2µg/ml) (cosmobio) using Lipofectamine3000 (Thermo). 48 hours post-transfection, cells were evaluated either by immunostaining or the insoluble fraction assay.

### Insoluble fraction assay

Insoluble fraction analysis was performed as previously described with minor modifications (Oji et al., 2020) Cultured cells were washed with ice-cold PBS and lysed in Triton X-100 buffer (50 mM Tris, 150 mM NaCl, 1 mM EDTA, 1% Triton X-100, 10% glycerol, and protease inhibitor cocktail (Thermo)). Lysates were then centrifuged at 100,000 x g for 20 min at 4°C. The supernatants were designated as the detergent-soluble fraction. Pellets were washed with Triton X-100 buffer, resuspended in SDS buffer (60 mM Tris-Cl (pH 6.8), 1 mM EDTA, 10% Glycerol, 2% SDS, and protease inhibitor cocktail (Thermo)), and sonicated for 5 s × 4 with a microtip sonicator. After centrifugation at 13,000 rpm for 15 min, the supernatants were identified as the detergent-insoluble fraction. Equal volumes of both insoluble and soluble fractions were boiled at 95°C for 10 min in SDS sample buffer and then analyzed by immunoblotting.

### siRNA transfection

Transfection of SH-SY5Y cells with siRNAs was carried out using Lipofectamine RNAiMax (Thermo Fisher Scientific) following the manufacturer’s instructions. The siRNAs used were detailed in Sasazawa et al., 2022, and are as follows: TRPML1 (SASI_Hs01_00067195), ALG2 (SASI_Hs01_00030269), Rab7a (SASI_Hs01_00104360), JIP4 (SASI_Hs01_00194613), TMEM55B (SASI_Hs02_00322347), and RILP CaMK2G (SASI_Hs01_00118118) from Sigma Aldrich. Additionally, non-coding siRNA was obtained from Dharmacon (Lafayette, CO).

### Ultrastructural analysis of SH-SY5Y cells by electron microscopy

Cells were first treated with a solution containing 2% glutaraldehyde (TAAB) and 50 mM sucrose (Wako) in a 0.1 M phosphate buffer at pH 7.4. This was followed by post-fixation with 1% osmium tetroxide in the same buffer. The fixed samples were then progressively dehydrated using a graded ethanol series and embedded in Epok812 (Okenshoji). Ultrathin sections, approximately 70 nm thick, were prepared using a UC6 ultramicrotome (Leica) and subsequently stained with uranyl acetate and lead citrate. The analysis was conducted using a transmission electron microscope (JEM-1400, JEOL).

## Acknowledgements

We thank Motoki Date for the Image J macro programing.

This work was supported by JST SPRING (Grant Numbers JPMJSP2109 to Y.D.), JSPS KAKENHI (Grant Numbers 18K15464, 21K07425, 18KK0242, 18KT0027 and 22H02986 to S.S.).

## Figure Legends

**Fig. S1.**
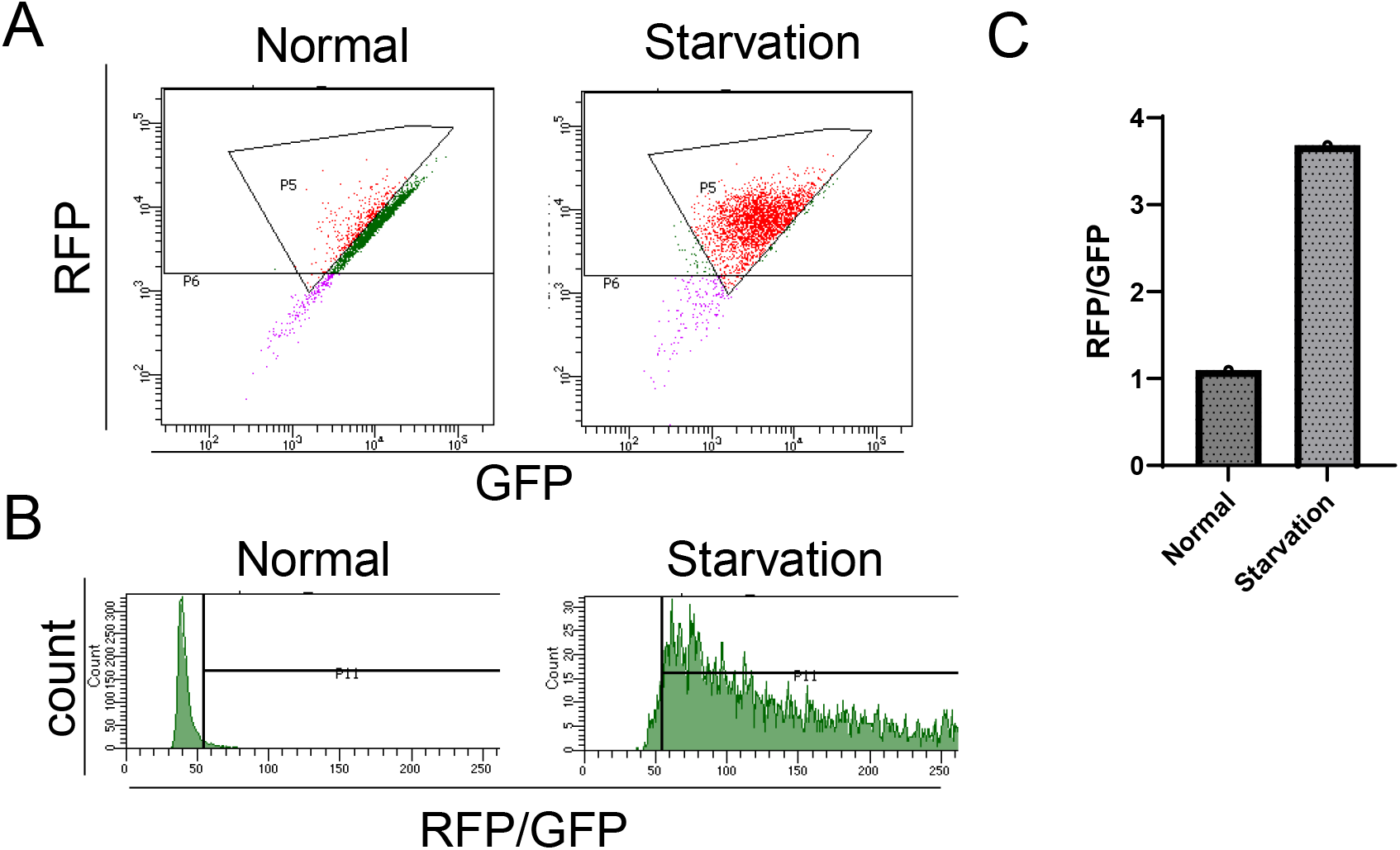
Flow cytometer analysis of RFP-GFP-LC3 during second screening. A-C. SH-SY5Y tetracycline-on (Tet-on) cells expressing RFP-GFP-LC3 were cultured in the presence of doxycycline (Dox). After Dox removal, the cells were treated with either normal medium or subjected to starvation for 24 h and then analyzed by flow cytometry. Dot plots display the gate-positive cell population (Autophagy induced cells: red). The histogram represents the number of cells on the vertical axis and the RFP/GFP ratio on the horizontal axis. The bar graph indicates the RFP/GFP ratio relative to the control.

**Fig. S2.**
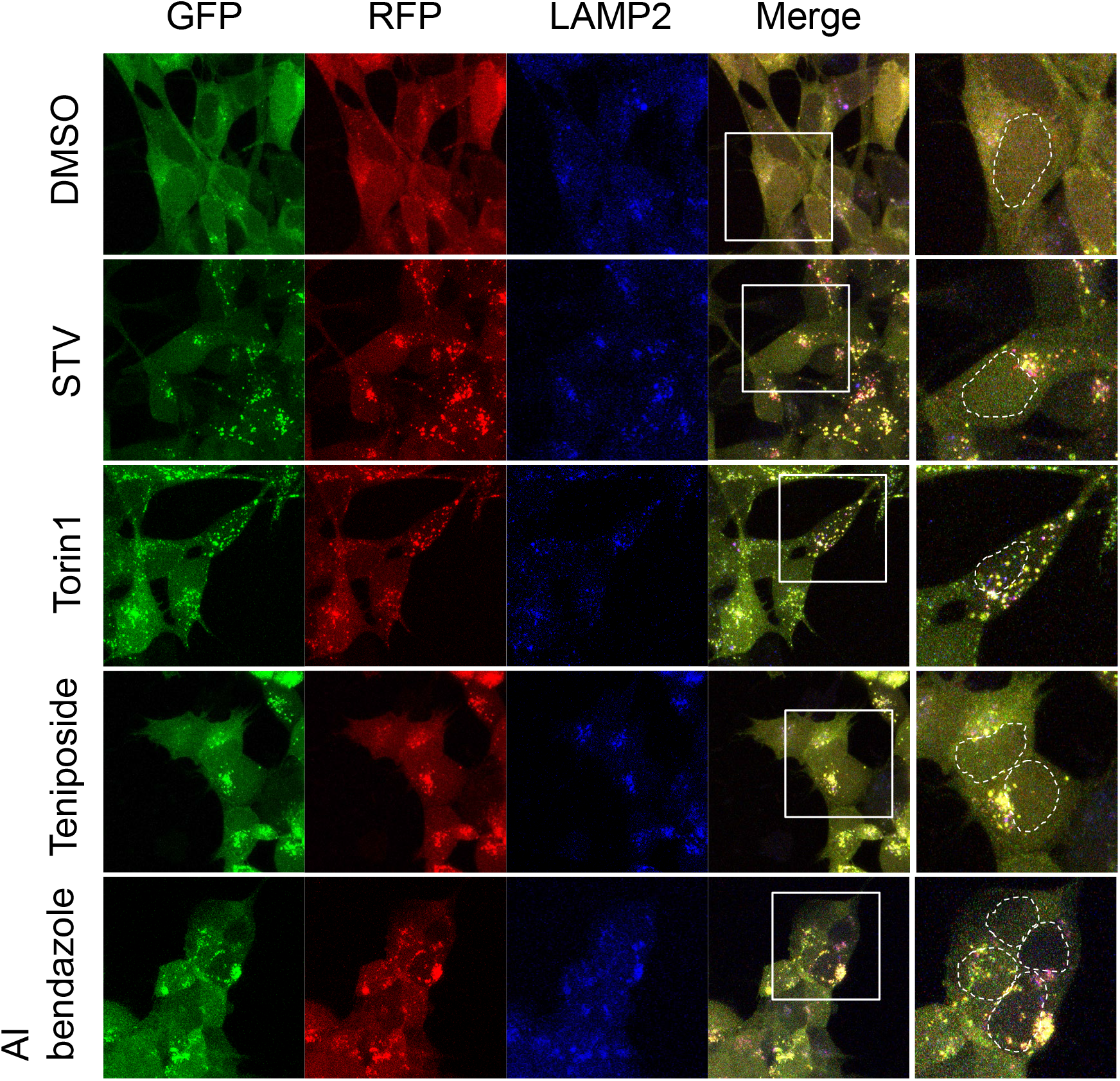
Confocal images of SH-SY5Y expressing RFP-GFP-LC3. SH-SY5Y cells expressing RFP-GFP-LC3 were treated with DMSO, starvation medium, torin1 (1 μM), teniposide (10 μM), or albendazole (10 μM). Post-treatment, cells were fixed and stained with an anti-LAMP2 antibody (blue: 647nm) and then imaged using a confocal microscope.

**Fig. S3.**
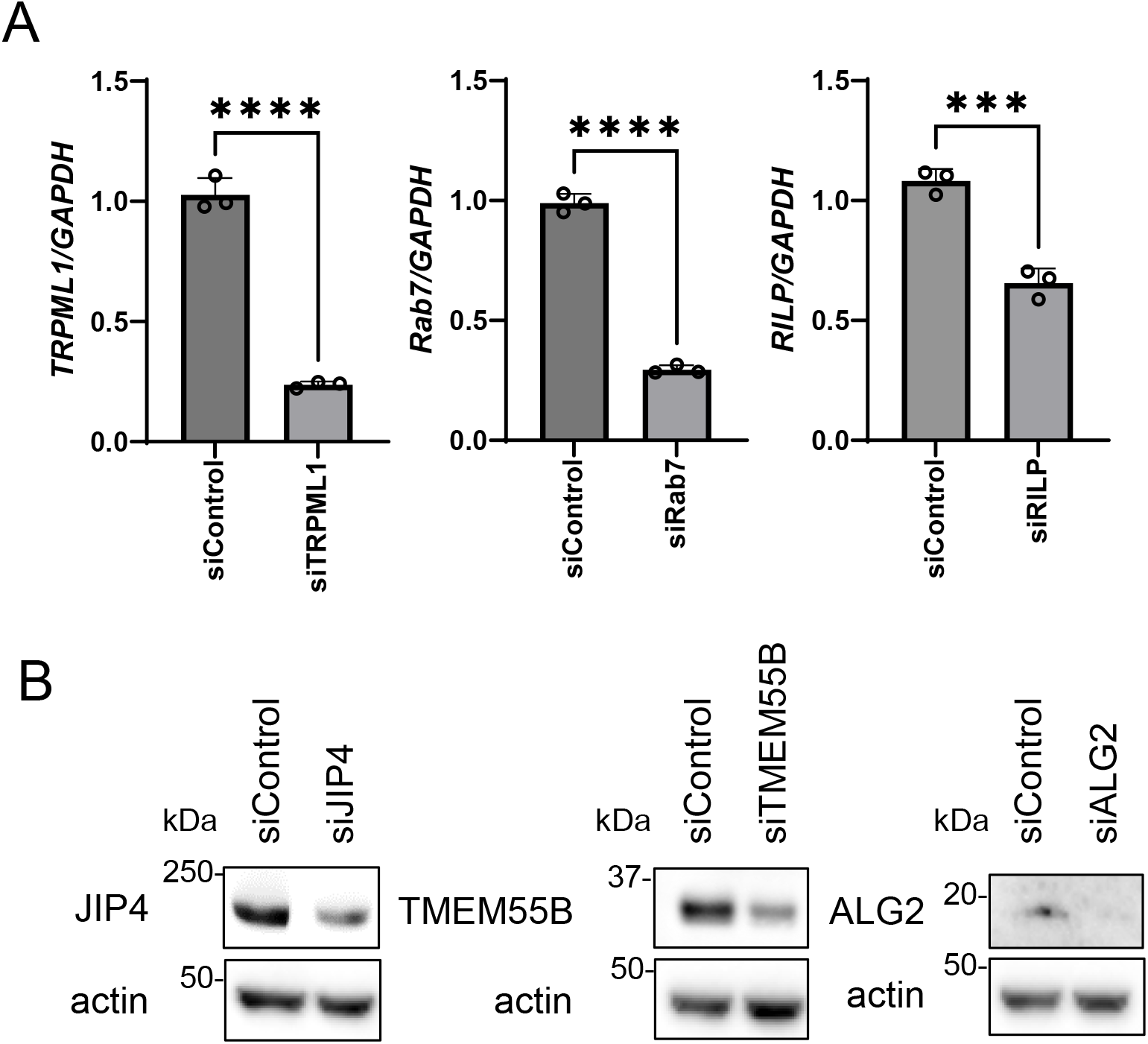
siRNAs and knockdown efficiency of each lysosomal factor. A, B. SH-SY5Y cells were treated with the specified siRNAs, and the knockdown efficiency of each siRNA was validated via qRT-PCR (n = 3 technical replicates) or Western blotting. The graph data is shown as the mean ± SD. ****P < 0.0001, ***P < 0.001; Student’s t-test.

**Fig. S4.**
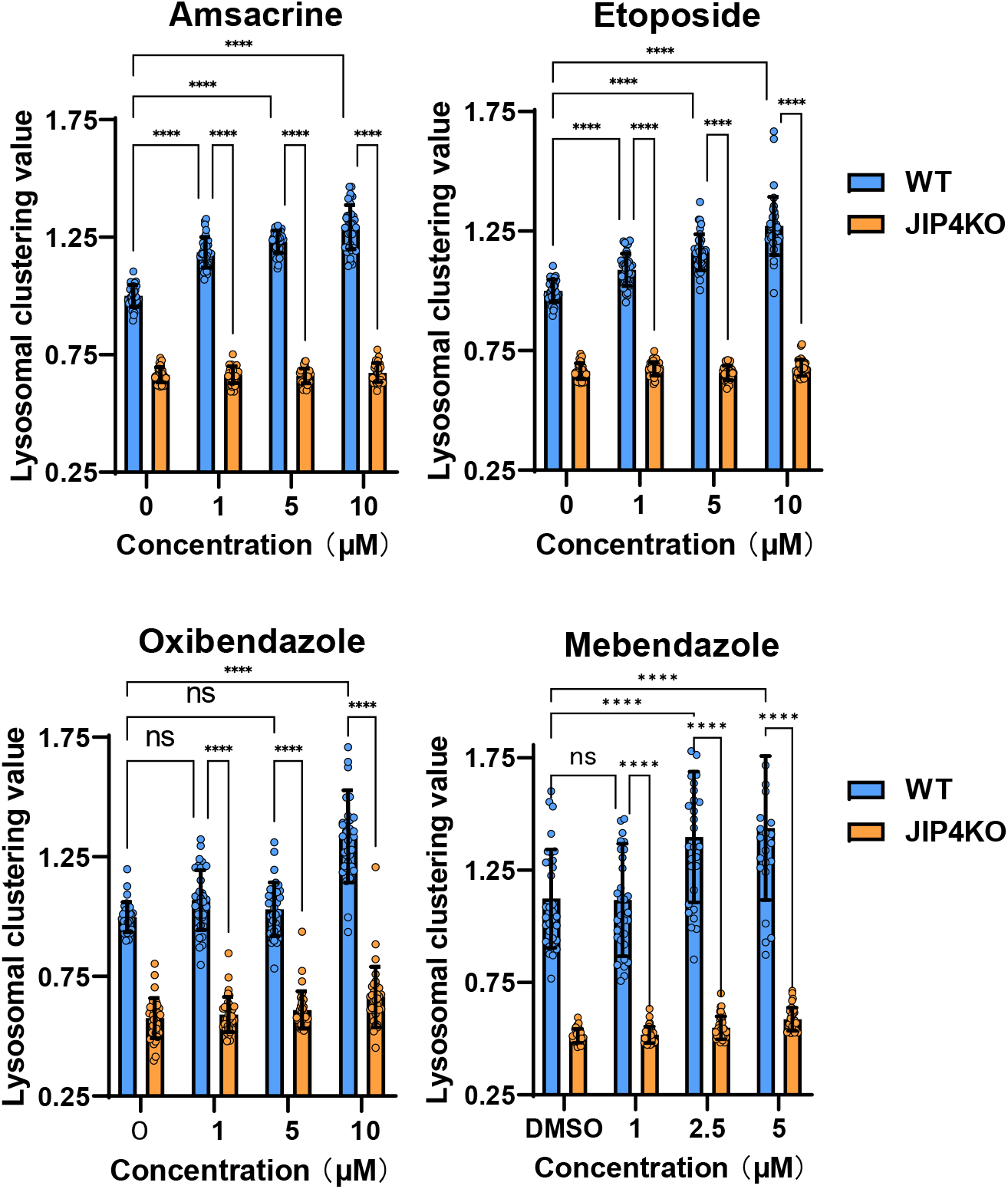
Lysosomal clustering analysis of other lysosomal clustering chemicals using JIP4KO cells. SH-SY5Y WT and JIP4 knockout cells were cultured in 96-well black plates and treated with amsacrine, etoposide, oxibendazole, or mebendazole at the specified concentrations. The cells were then fixed and stained with anti-γ-tubulin (green) and an anti-LAMP2 antibody (red) before imaging with INCellAnalyzer2200. INCellAnalyzer2200 images were processed and analyzed using ImageJ for lysosomal clustering. The graph displays the lysosomal clustering values of either WT or JIP4KO cells. (n > 30). Data is represented as the mean ± SD. ****P < 0.0001, ***P < 0.001, **P < 0.01, *P < 0.05; N.S. denotes not statistically significant, determined via two-way ANOVA and Tukey’s test.

**Fig. S5.**
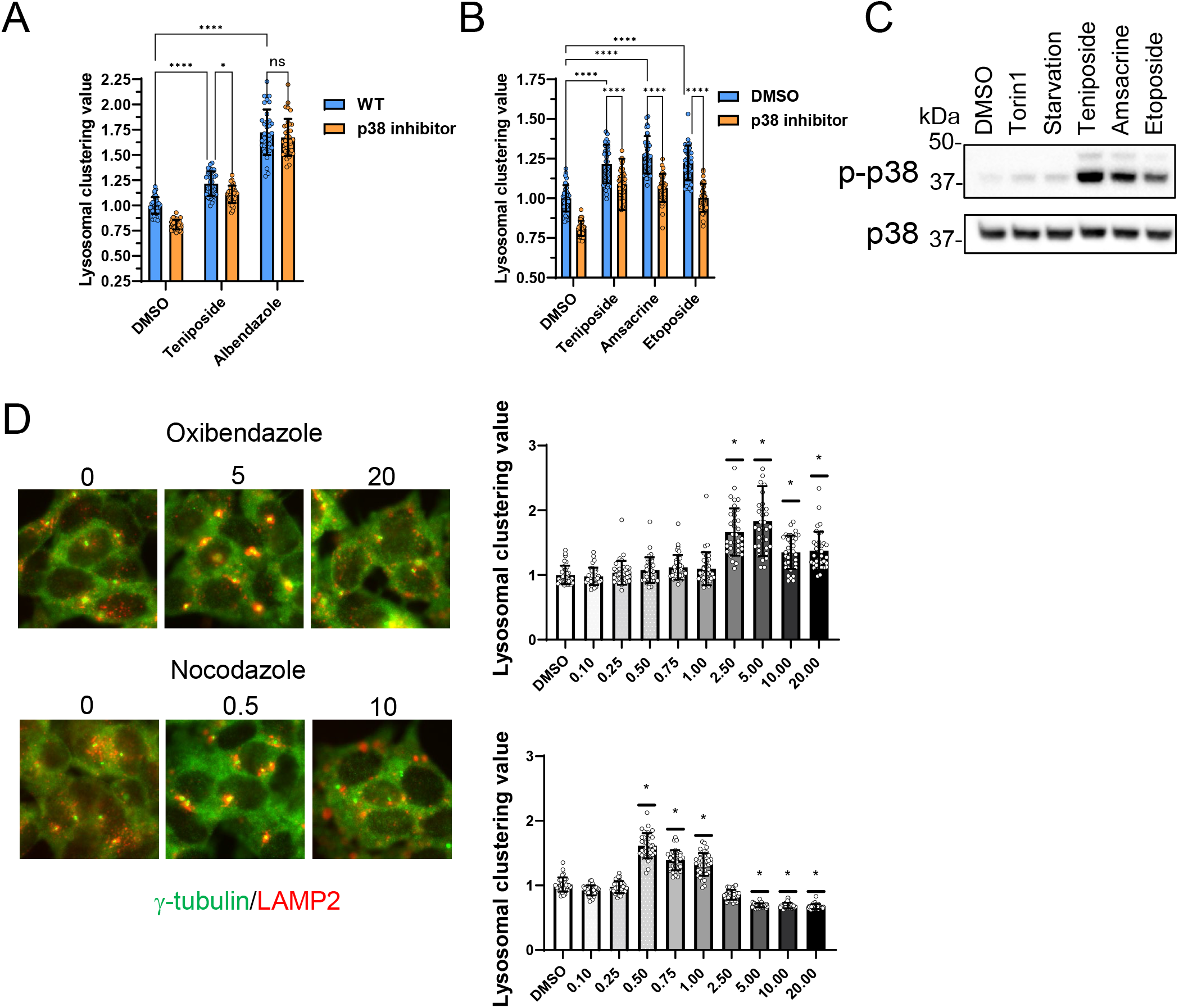
Analysis of lysosomal clustering mechanism of action for topoisomerase inhibitors and the benzimidazole mechanism. A, B. SH-SY5Y cells were cultured in 96-well black plates and treated with teniposide (10 μM), amsacrine (10 μM), etoposide (10 μM), or albendazole (10 μM) with or without the p38 inhibitor (SB203580). Following treatment, cells were fixed and stained with anti-γ-tubulin (green) and an anti-LAMP2 antibody (red). They were then imaged using INCellAnalyzer2200. INCellAnalyzer2200 images were analyzed for lysosomal clustering with ImageJ. The graph presents the lysosomal clustering value for either DMSO or the p38 inhibitor. (n > 30). Data is shown as the mean ± SD. ****P < 0.0001, ***P < 0.001, **P < 0.01, *P < 0.05; N.S. denotes not statistically significant, determined by two-way ANOVA and Tukey’s test. C. SH-SY5Y cells were cultured and then treated with starvation medium, torin1 (1 μM), teniposide (10 μM), amsacrine (10 μM), or etoposide (10 μM). Subsequently, cell lysates were immunoblotted with the indicated antibodies. D. SH-SY5Y cells were cultured in 96-well black plates and treated with Oxibendazole or Nocodazole at specified concentrations (µM). After treatment, cells were fixed and stained with anti-γ-tubulin (green) and an anti-LAMP2 antibody (red), followed by imaging with INCellAnalyzer2200. INCellAnalyzer2200 images were processed and analyzed using ImageJ for lysosomal clustering. The graph displays the lysosomal clustering values. (n > 30). Data is presented as the mean ± SD. *P < 0.0001, as determined by one-way ANOVA and Dunnet’s test (vs. DMSO).

